# DNA satellite and chromatin organization at house mouse centromeres and pericentromeres

**DOI:** 10.1101/2023.07.18.549612

**Authors:** Jenika Packiaraj, Jitendra Thakur

**Affiliations:** Department of Biology, Emory University, 1510 Clifton Rd, Atlanta, GA 30322

**Keywords:** CENP-A, H3K9me3, constitutive heterochromatin, long-read sequencing, transposable elements, repetitive DNA

## Abstract

Centromeres are essential for faithful chromosome segregation during mitosis and meiosis. However, the organization of satellite DNA and chromatin at mouse centromeres and pericentromeres is poorly understood due to the challenges of sequencing and assembling repetitive genomic regions. Using recently available PacBio long-read sequencing data from the C57BL/6 strain and chromatin profiling, we found that contrary to the previous reports of their highly homogeneous nature, centromeric and pericentromeric satellites display varied sequences and organization. We find that both centromeric minor satellites and pericentromeric major satellites exhibited sequence variations within and between arrays. While most arrays are continuous, a significant fraction is interspersed with non-satellite sequences, including transposable elements. Additionally, we investigated CENP-A and H3K9me3 chromatin organization at centromeres and pericentromeres using Chromatin immunoprecipitation sequencing (ChIP-seq). We found that the occupancy of CENP-A and H3K9me3 chromatin at centromeric and pericentric regions, respectively, is associated with increased sequence abundance and homogeneity at these regions. Furthermore, the transposable elements at centromeric regions are not part of functional centromeres as they lack CENP-A enrichment. Finally, we found that while H3K9me3 nucleosomes display a well-phased organization on major satellite arrays, CENP-A nucleosomes on minor satellite arrays lack phased organization. Interestingly, the homogeneous class of major satellites phase CENP-A and H3K27me3 nucleosomes as well, indicating that the nucleosome phasing is an inherent property of homogeneous major satellites. Overall, our findings reveal that house mouse centromeres and pericentromeres, which were previously thought to be highly homogenous, display significant diversity in satellite sequence, organization, and chromatin structure.

## INTRODUCTION

Centromeres are the chromosomal sites where spindle fibers attach via the kinetochore to allow chromosome segregation during cell division. Defects in centromere function can cause chromosome missegregation and aneuploidy, which are linked to cancers, miscarriages, and genetic disorders [1-4]. Centromeres are characterized by specialized nucleosomes composed of Centromere-Protein A (CENP-A), which replaces canonical histone H3 at centromeric chromatin [5, 6]. CENP-A chromatin acts as the foundation for the assembly of kinetochore components. In mammals, CENP-A is assembled on long arrays of tandem DNA repeats called satellites [7]. Human centromeres comprise α-satellite (171 bp monomer) arrays, some of which are further organized in higher-order repeat structures [8-10]. A highly homogeneous α-satellite core forms the functional centromere, which is flanked by more divergent α-satellite monomers [8-11]. The human pericentromeric regions comprise several different types of complex sequences, including other satellites, genes, and transposable elements (TEs) [9, 12]. TEs are also found at the functional centromeres of other eukaryotes, including *Drosophila* [13]. Centromeric satellite sequences and organization can vary greatly, even between chromosomes within the same individual, as seen in humans [9, 14-18].

Due to the lack of conserved centromeric sequences, CENP-A chromatin is considered the epigenetic mark of centromeres. This is further supported by the formation of functional ectopic centromeres, called neocentromeres, at locations lacking satellite sequences [19-21]. CENP-A chromatin has been extensively studied through in vitro reconstitution, demonstrating the presence of octameric, hexameric, and tetrameric CENP-A nucleosomes in various eukaryotes [22-27]. In vivo studies using tagged CENP-A pulldown have also revealed the existence of CENP-A dimers within nucleosomes. However, the centromeric chromatin organization on satellite arrays *in vivo* remains poorly understood. Studies in humans suggest that a 340 bp α-satellite dimeric unit is occupied by two CENP-A particles bridged by a CENP-B, CENP-C, and CENP-T containing linker [28, 29]. Furthermore, sequence variations across different α-satellite dimers within a given array on a given chromosome corresponded to variations in CENP-A chromatin profiles, suggesting a sequence-dependent assembly of centromeric chromatin [30]. Pericentromeric regions assemble distinct constitutive heterochromatin in which histone H3 is trimethylated at its lysine 9 residue (H3K9me3) [31-33]. Pericentric heterochromatin binds to cohesin, which is required for proper chromosome segregation by preventing sister chromatid separation before anaphase [34, 35].

Sequencing and assembling centromeres have been challenging due to the highly repetitive nature of centromeric DNA [36]. As a result, centromeres and other repetitive elements have been omitted or only partially annotated in genome assemblies. The lack of centromere assemblies has thus limited studies of centromeric chromatin structure using genomics-based chromatin profiling methods. However, recent advances in high-fidelity long-read sequencing (LRS) have opened the possibility for further in-depth analysis of centromere organization and chromatin structure [37, 38]. In addition, the LRS technologies have led to the development of the Telomere-to-Telomere (T2T) gapless human genome assembly, which has allowed the characterization of centromeric and pericentromeric arrays in humans [9, 39].

Unlike human centromere satellites that have been extensively characterized with the advent of advanced genomics technologies, studies to comprehensively characterize mouse centromeric satellite arrays have begun only recently. Mouse centromeres are defined by arrays of minor satellites (120 bp monomer) [40, 41]. Minor satellite (MiSat) arrays are flanked by pericentromeric major satellites (MaSat) (234 bp monomer) [42]. MiSats are associated with the centromere proteins such as CENP-A, CENP-B, and CENP-C, while MaSats are associated with heterochromatin protein 1α (HP1) [43-45]. Both MaSat and to a lesser extent, MiSat, have been shown to contain H3K9me3 [43, 45]. H3K9me3 is shown to exhibit a specific repeating dinucleosomal configuration on major satellites, while minor satellites display simple mononucleosomal H3K9me3 configuration [43]. Unlike human α-satellites, which share 60-100% sequence similarity, mouse MiSat and MaSat arrays were previously thought to be highly homogeneous with few sequence variations within an array and between chromosomes [46, 47]. Recent studies have identified a considerable sequence heterogeneity and copy number of variations of MiSat across different mouse populations and strains [44, 48]. The variations include sites of high sequence variation at the 17 bp CENP-B box motif that binds Centromere Protein B (CENP-B) [48]. Furthermore, analyses of Sanger sequencing traces from C57BL/6 (B6) have shown that continuous MiSats are rarely identical and are likely to vary at the CENP-B box [49]. It remains unclear how MiSats and MaSats are arranged across long regions at centromeres and pericentromeres.

In this study, we investigated the organization of MiSat and MaSat arrays and associated chromatin in *Mus musculus* reference strain C57BL/6 (B6). First, we identified long satellite arrays by analyzing publicly available PacBio LRS data [50]. We found variations in both sequence and organization of MiSats and MaSats within and across arrays in this genome. We found a surprisingly high level of variation in the CENP-B box sequence within MiSat arrays. Although the majority of satellites were present as continuous arrays, we also detected TEs interspersed with satellites in a significant fraction of both MiSat and MaSat arrays. Subsequently, we analyzed the organization of CENP-A chromatin along with constitutive H3K9me3 and facultative H3K27me3 heterochromatin at centromeric MiSat and pericentromeric MaSat arrays by generating high-resolution Chromatin immunoprecipitation Sequencing (ChIP-seq) data for CENP-A, H3K9me3, and H3K27me3. We found that the enrichment of CENP-A and H3K9me3 at both centromeric and pericentromeric regions differs on arrays containing different satellite variants. Furthermore, TEs at centromeric regions were not bound to CENP-A, indicating their absence from the functional centromeric domains. Finally, we found that while homogeneous MaSat arrays contain H3K9me3 nucleosomes in a well-phased configuration, homogeneous MiSat arrays contain CENP-A nucleosomes that lack a phased configuration.

## RESULTS

### Mouse MiSats and MaSats are organized as continuous and interspersed satellite arrays

We analyzed publicly available LRS data from C56BL/6J mouse strain generated using PacBio Sequel II System with HiFi sequencing, which yields highly accurate (99.8%) long reads [50]. Using NCBI-BLAST, we identified MiSat and MaSat arrays in the LRS data using reference MiSat and MaSat consensus units as query sequences. The majority of long reads containing MiSat and MaSat ranged from 13 kb to 19 kb in length (Supplementary Figure 1A). Analysis of the LRS reads using RepeatMasker revealed that MiSat and MaSat arrays also contain a significant amount (4.56% in MiSat reads and 11.26% in MaSat reads) of non-satellite sequences (Figure 1A). These non-satellite sequences include repeats, such as transposable elements, simple repeats, and other unknown sequences. Next, we analyzed the arrangement of satellite and non-satellite sequences on arrays. Satellite containing long reads displayed two distinct organizations: Type 1 continuous arrays (98.23% of MiSat arrays and 90.05% of MaSat arrays) and Type 2 arrays interspersed with non-satellite sequences (1.77% of MiSat arrays and 10% of MaSat arrays) (Figure 1B, Supplementary Figure 1). Type 1 continuous MiSat and MaSat arrays included a subset of arrays (1.92% of MiSat arrays and 0.85% of MaSat arrays) where monomers switched direction from forward to reverse or vice-versa (Figure 1B-1C). Sequences interspersed within Type 2 MiSat arrays predominantly comprised Long Terminal Repeat (LTR) retrotransposons (present in 56.38% of Type 2 MiSat arrays) (Figure 1B -1E) with the IAPEz-int family, part of the intracisternal A-type particle (IAP) class of endogenous retroviruses (ERV2) being the most abundant (Figure 1E).

**Figure 1.**
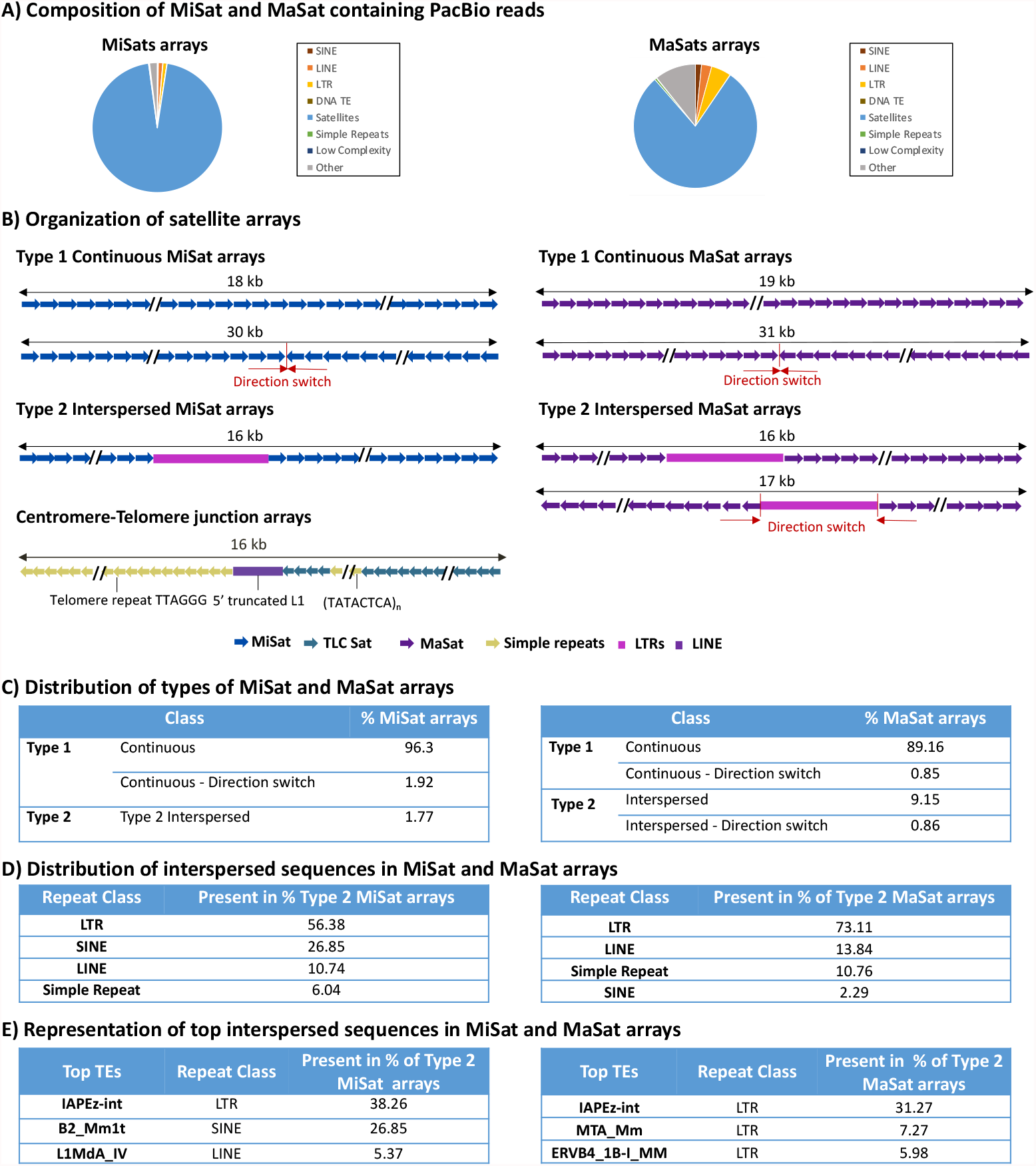
Variations in MiSat and MaSat array composition and organization. **A)** Composition of reads containing MiSats (Left) and MaSats (Right). While MiSats and MaSats are the most abundant sequences, non-satellite sequences constitute a significant proportion of these reads. **B)** Organization of representative MiSat and MaSat arrays seen as Type 1 continuous arrays and Type 2 arrays interspersed with non-satellite sequences. Additionally, satellite organization at a representative Centromere-Telomere junction is shown. **C)** Distribution of Types of MiSat (Left) and MaSat (Right) arrays **D)** Distribution of interspersed sequences in Type 2 MiSat (Left) and MaSat (right) arrays **E)** Representation of top interspersed sequences in Type 2 arrays. These top interspersed sequences are the most frequently occurring non-satellite sequences within the Type 2 MiSat and MaSat arrays.

The IAPEz-int family contains young TEs that have been studied for their roles as functional transcriptional promoters of nearby genes and epigenetic modulators through DNA methylation and H3K9 modifications [51-53]. Another abundant TE interspersed with Type 2 MiSat arrays was the B2 element, which belongs to the Short interspersed nuclear element (SINE) class of non-LTR retrotransposons (Figure 1E). B2 elements in mice have been shown to be present at boundaries between H3K9me3 and H3K9me2 chromatin domains [54] and provide CCCTC-binding factor (CTCF) binding sites [55, 56]. Similarly, Type 2 MaSat arrays are mostly interrupted by LTR retrotransposons (31.27%), including those from the IAPEz-int family (ERV2), MTA (ERV3), ERVB4_1B (ERV2), RLTR6 family (ERV1), RLTR10 family (ERV2), and MERVL family (ERV3).

Next, we investigated the satellite organization at the junctions of centromeres and telomeres. Telomere and centromere junctions had a distinct organization with four types of sequences (Figure 1D): TeLoCentric (TLC) satellites, a short stretch of (TATACTCA)_n_ simple repeats, 5’ truncated L1 element, and telomeric repeats (TTAGGG)_n_. TLC satellites are 145-146 bp repeats found near telomeres in most *Mus musculus* species that share 60-70% sequence homology with minor satellites [46]. The 5’ truncated LINE-1/L1 is a previously reported highly conserved element of centromere telomere junctions [46]. L1 is part of the Long interspersed nuclear elements (LINE) group of non-LTR retrotransposons that is highly abundant in almost all mammalian genomes [57].

### Mouse MiSats and MaSats exhibit high sequence variations within and across different arrays

To investigate the sequence similarity among repeat units within and across satellite arrays, we compared and aligned satellite monomers isolated from a given LRS read with the *M. musculus* reference MiSat and MaSat satellite units. Similar to human α-satellites, mouse MiSats contain a 17 bp sequence motif called the CENP-B box that binds to CENP-B centromeric protein in a sequence-dependent manner [58-60]. CENP-B is the only centromeric protein that binds to its target satellite sequences in a sequence-dependent manner. Although CENP-B was initially thought to be dispensable for centromere function [61], recent studies have shown its critical role in the maintenance of centromeric memory [62]. Interestingly, we found that MiSat units in the B6 strain exhibited significant nucleotide variations from the reference (88.6-98.3%), with most changes concentrated at and around the 17 bp CENP-B box, especially at positions 15-17 (Figure 2A-C, Supplementary Figure 2). As a result, an intact CENP-B box was present only in a subset of satellite units in each array (Figure 2A). For Type 2 arrays with interspersed non-satellite sequences, sequence variation was present at either side of the interrupting non-satellite sequence (Figure 2A). In addition to the sequence variations within arrays, we found striking variations in organization between different types of arrays. A subset of MiSat arrays comprises divergent monomers with different monomer lengths: 112-mer (7.78%) and 112-64-dimer (4.56%), which were previously reported by Rice (2020) (Figure 2B). The density of intact CENP-B Boxes varied greatly between variant 1 112-mer arrays and 112-64-dimer arrays. Type 1 112-mer arrays contained a few intact CENP-B Boxes, while Type 1 112-64-dimer arrays contained a high number of intact CENP-B Boxes (Figure 2B, 2E). Furthermore, TLC satellite arrays, which lack CENP-B boxes, displayed high nucleotide variation (56.1-95.2%) within the same array (Figure 2D).

**Figure 2.**
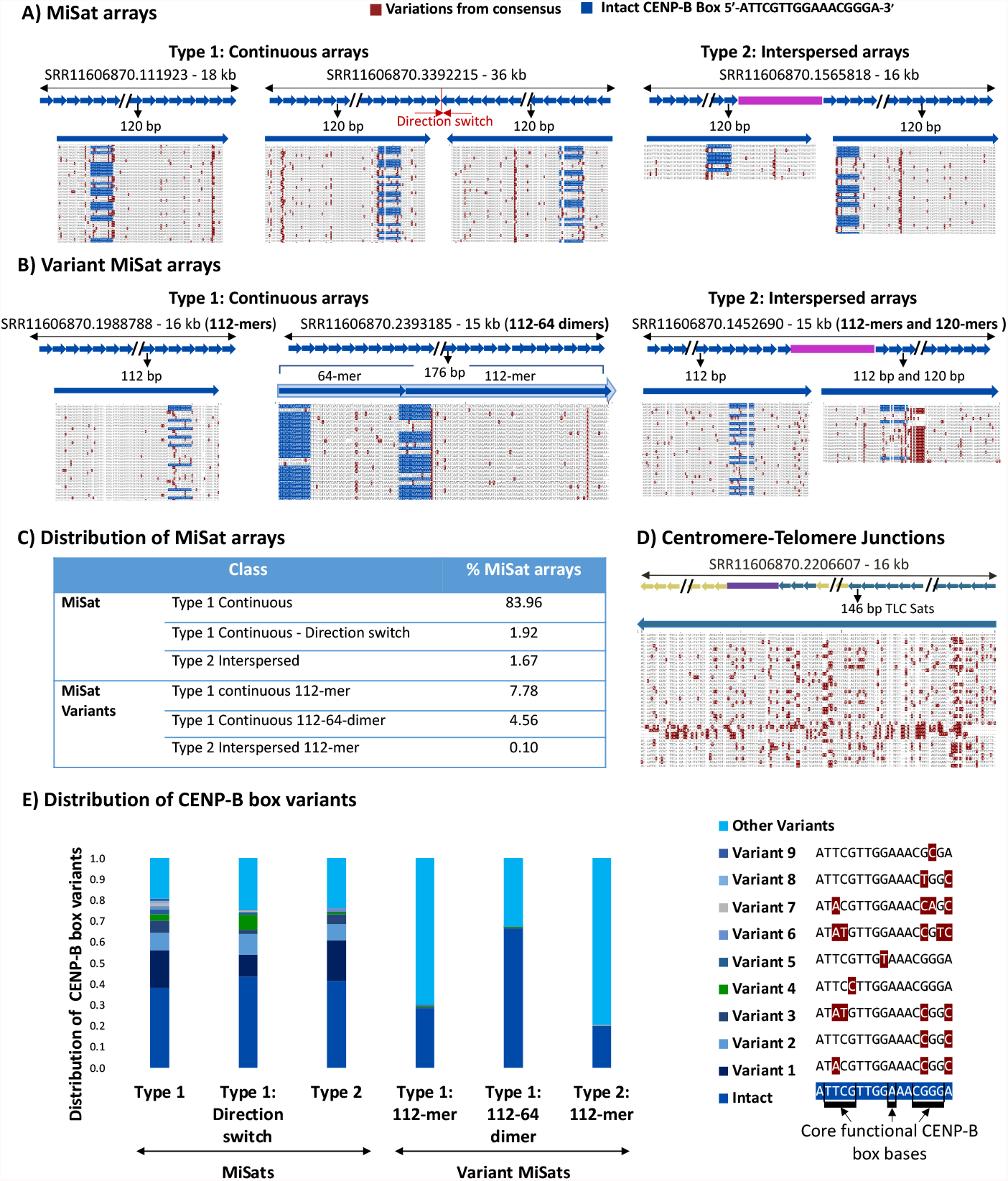
Sequence composition of satellite units within and across centromeric MiSat arrays. Alignments of **A)** the MiSat consensus with repeat units from representative MiSat arrays and, **B)** the 112-mer and 112-64-dimer MiSat consensus with repeat units from representative variant MiSat arrays. **C)** The distribution of different types of MiSat arrays. **D)** Alignment of the TLC satellite consensus with repeat units from representative arrays. Reference satellite consensus sequences used for the alignments are as described previously [47, 49, 63]. Alignments of repeat units from the beginning of a given array are shown. Alignments of all the repeat units from arrays are given in Supplementary Figure 2. **E)** The distribution of CENP-B box variants in different types of MiSat arrays.

Similarly, we found satellite variations in MaSat-containing arrays (Figure 3A-C, Supplementary Figure 3). While most Type 1 and Type 2 MaSat arrays contained highly homogeneous monomers with a low level of sequence variation within a single array (93.7-98.7), including at MaSat motif 5’-GAAAACTGAAAA -3’ (Figure 3A, 3C). However, a subset of Type 1 and Type 2 major satellite arrays (10.01%) contained diverged monomers that exhibited increased nucleotide variation from the consensus (65.3-79.9%) (Figure 3B, 3C). Nucleotide variations in divergent MaSat arrays included several insertions and deletions, leading to variations in major satellite monomer lengths such as 220-mers and 250-mers (Figure 3B).

**Figure 3.**
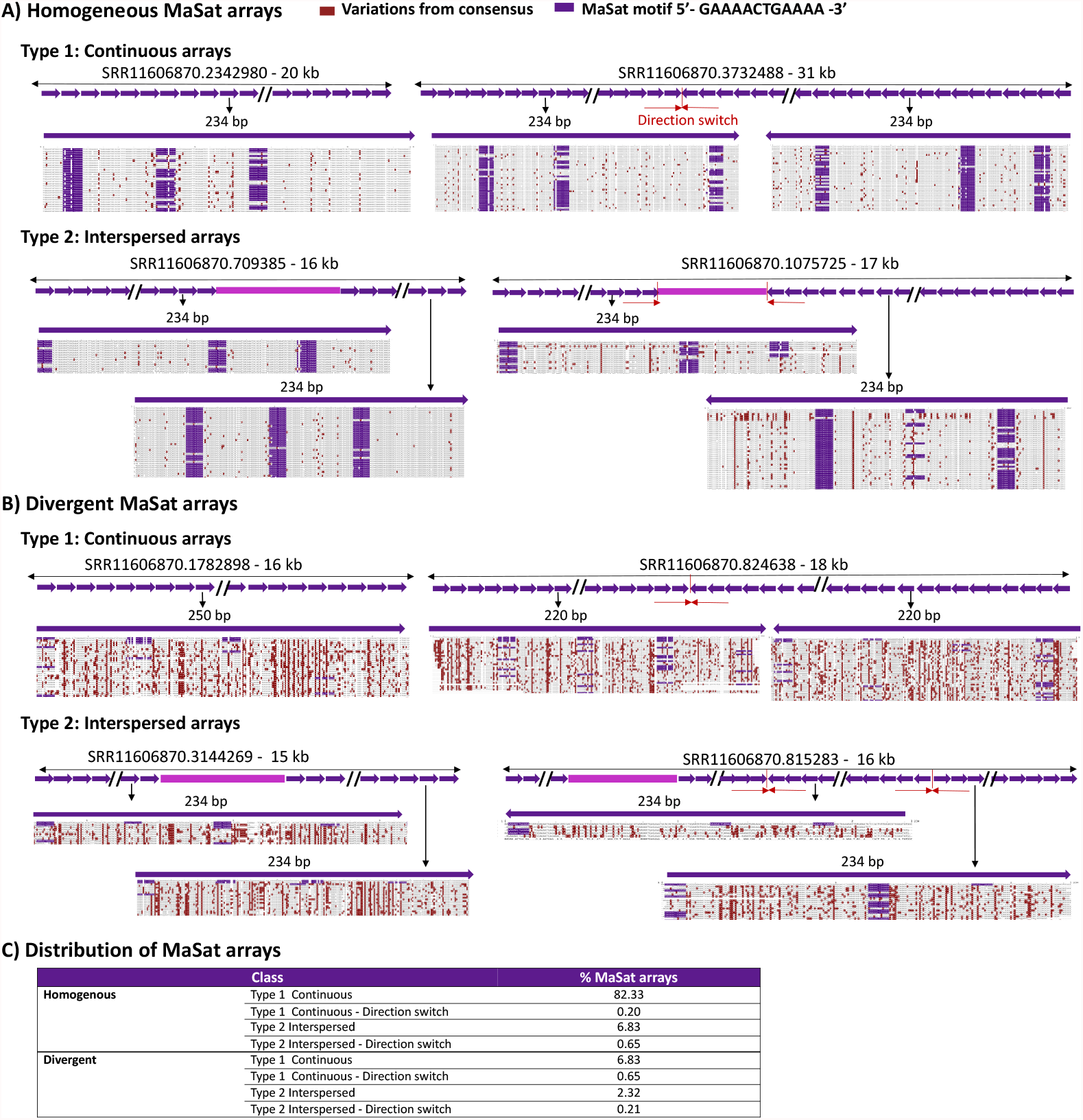
Sequence composition of satellite units within and across pericentromeric MaSat arrays. Alignments of **A)** the MaSat consensus with repeat units from four representative MaSat homogeneous arrays, and **B)** the MaSat consensus sequences with repeat units from four representative divergent arrays. Reference satellite consensus sequences used for the alignments are as described previously [63]. Alignments of repeat units from the first part of a given array are shown. Alignments of the repeat units from full arrays are given in Supplementary Figure 3. **C)** The distribution of different types of MaSat arrays.

Next, to investigate the arrangement of MiSat and MaSat arrays on longer contigs, we generated assembled contigs from the PacBio reads using Hifiasm [64]. We classified resulting contigs containing MiSats into five type based on their organizational patterns (Figure 4A). The first class, which was the most abundant, included MiSat arrays transitioning to telomeres, encompassing TLC Sats, 5’ truncated L1, and simple repeats. Interestingly, all these contigs contained a direction change near the MiSat to TLC Sat transition. The length of the TLC sat arrays within these contigs ranged from 44-50 TLC monomers. A few contigs in this class contained interspersed IAPEz-int TEs. The second class of contigs solely contained long continuous MiSat arrays. The third class of contigs displayed a sharp transition from MaSats to MiSats. The fourth class included two contigs that contained interspersed 112-mer and 120-mer MiSat transitioning to MaSat. The fifth class featured a 1Mb long contig encompassing MaSat, MiSat, TLC sat, and telomere repeats. This contig contained 112-64-mer dimer MiSat variant at the transition from MiSat to MaSat and IAPEz-int interspersed within the MiSat. Similar to MiSat contigs, we classified contigs containing MaSats into five types based on their organizational patterns (Figure 4B). The first class contained long continuous homogeneous MaSat contigs. The second class contained homogeneous MaSats interspersed with TEs and simple repeats. The third class included contigs that contained both homogeneous and MaSat and divergent MaSat. Contigs were classified as having MaSats and divergent MaSats if over 20% of the monomers in the same contig were divergent. The fourth class encompassed MaSats, divergent MaSats, IAPEz-int TEs, and simple repeats with or without a direction change. The fifth class included contigs in which MaSats transitioned into long arrays of simple repeats, TEs, and rDNA. Overall, the composition and arrangement pattern of MiSat and MaSat arrays we defined using unassembled PacBio reads (Figure 1-3) were captured in the contigs assembled from these reads using the Hifiasm assembler.

**Figure 4.**
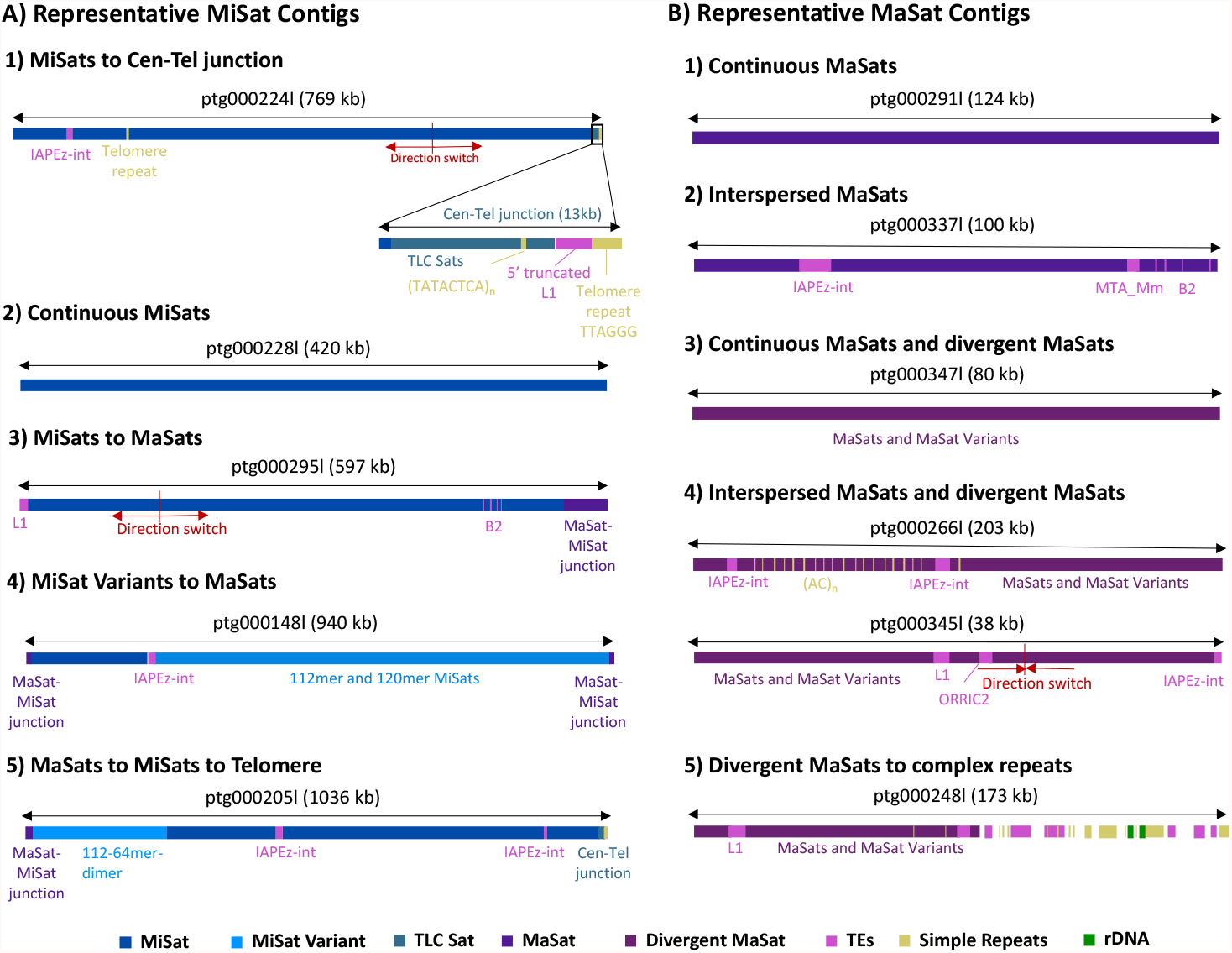
Organization of MiSat and MaSat arrays on assembled contigs. MiSat and MaSat contigs obtained through the Hifiasm assembler were classified into different types based on distinct organizational patterns. **A)** MiSat contigs types 1) Include MiSats transitioning to telomeres, containing TLC Sats, 5’ truncated L1, and simple repeats. These contigs exhibit a change in direction near the MiSat to TLC Sat transition and occasionally contain interspersed TEs such as IAPEz-int. 2) Include long uninterrupted MiSat arrays. 3) Include MiSats transitioning to MaSat with a sharp boundary. 4) Include interspersed MiSat variants transitioning to MaSat. 5) Include MaSat, MiSat, MiSat variants, IAPEz-int, TLC sat, and telomere repeats. **B)** MaSat contig types 1) Include long uninterrupted MaSat contigs. 2) Include MaSat interspersed with TEs and simple repeats. 3) Include MaSats and MaSats variants 4) Include MaSats, MaSat variants and interspersed TEs with and without a direction change 5) MaSats transition into complex repeats, such as simple repeats, TEs, and rDNA.

### Abundant 120-mer Misat arrays are preferred as functional centromeres

To determine if MiSat array types differ in chromatin assembled at mouse centromeric regions, we performed ChIP-seq for CENP-A, H3K9me3, and H3K27me3 in the B6 strain. We mapped the sequencing data to representative MiSat array types (Figure 5A) and calculated enrichment on each array type by normalizing ChIP enrichment with the abundance of the respective array in the ChIP input (Figure 5B). Among all MiSat types, we observed the highest CENP-A enrichment on abundant 120-mer Type 1 and Type 2 arrays. Within Type 2 interspersed MiSat arrays, CENP-A was enriched at MiSats but not at non-satellite regions, suggesting that TEs interrupting MiSat arrays are not part of functional kinetochores. However, while the IAPEz-int elements interrupting Type 2 MiSat were not enriched in CENP-A, they were significantly enriched in H3K9me3, suggesting that they are repressed at centromeres (Figure 5A-5B). At centromere and telomere junctions, TLC satellites were not enriched for CENP-A or H3K9me3 (Figure 5B, Supplementary Figure 4). However, the 5’ truncated L1 element present at these junctions showed H3K9me3 and H3K27me3 enrichment, indicating the heterochromatic nature of these sites (Figure 5B, Supplementary Figure 4). In general, divergent MiSat arrays showed a low level of CENP-A (Figure 5A-5B). Interestingly, divergent arrays containing 112-64-dimeric units also showed a significant CENP-A enrichment (Figure 5A-5B). Overall, abundant 120-mer Type1 continuous and Type 2 interspersed MiSat arrays are preferred as functional centromeres, as they exhibit high enrichment of the CENP-A, a chromatin mark that targets chromosomal loci for functional centromere formation. Furthermore, a significant enrichment of H3K9me3 on TE elements at centromeric regions indicates their heterochromatic and silenced nature.

**Figure 5.**
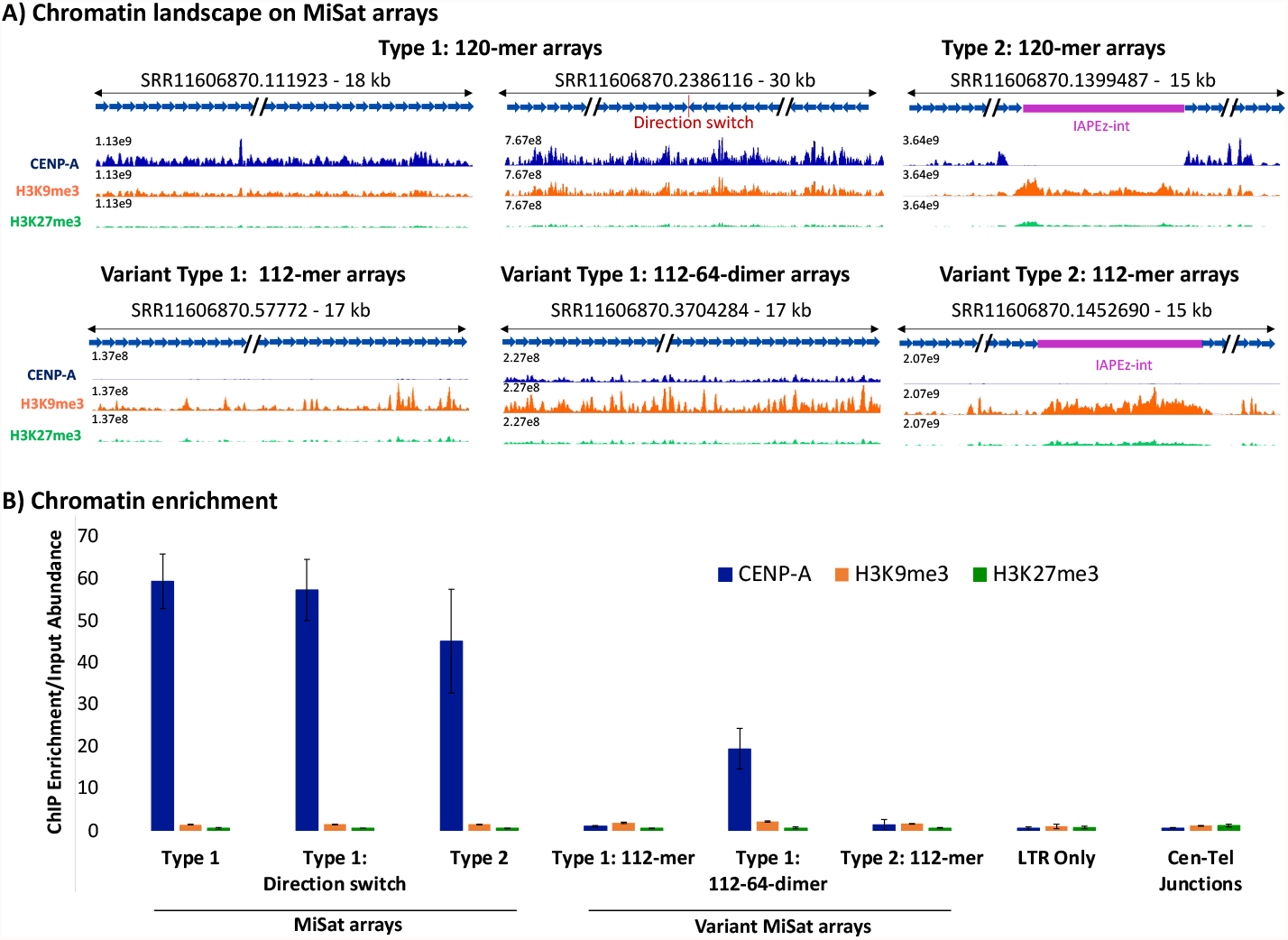
Occupancy of chromatin marks on MiSat arrays. **A)** CENP-A, H3K9me3, and H3K27me3 chromatin profiles on representative abundant 120-mer MiSat arrays, and variant 112-mer and 112-64-dimeric MiSat arrays. The Y-axis range is set to a constant value for a given array for CENP-A, H3K9me3, and H3K27me3 tracks. The length of each array is given, and the X-axis is not to the scale. **B)** CENP-A, H3K9me3, and H3K27me3 enrichment. Overall enrichment on each MiSat array type is calculated by normalizing the ChIP enrichment with the abundance of the respective MiSat array in the ChIP input. The normalized enrichment value was averaged over three or more arrays for each type.

### Homogeneous MaSat arrays exhibit increased constitutive heterochromatin at pericentric regions

To determine the enrichment of H3K9me3 chromatin at mouse pericentric regions, we mapped the sequencing data to representative MaSat arrays for both Type 1 and Type 2 arrays (Figure 6A). As expected, MaSat arrays were enriched for H3K9me3 (Figures 6A and 6B). Additionally, we found that in most type 2 MaSat arrays, the interrupting LTR transposon was enriched in H3K9me3. However, the H3K9me3 enrichment on MaSat was significantly higher than on the interrupting LTR transposon (Figures 6A and 6B). Furthermore, we found that divergent MaSat arrays were also enriched for H3K9me3, albeit at a ∼2-fold lower level compared to homogeneous MaSat arrays. Interestingly, divergent MaSat arrays were enriched for slightly higher amounts of H3K27me3 facultative heterochromatin as compared to their homogeneous counterparts (Figures 6A and 6B). In these divergent interspersed arrays, the H3K9me3 enrichment at the interrupting LTR transposon was much higher than the H3K9me3 enrichment at MaSat (Figure 6A). Together, these results suggest that sequence homogeneity within MaSat arrays is highly correlated with the presence of constitutive heterochromatin. As the sequence homogeneity decreases, MaSat arrays start assembling facultative heterochromatin.

**Figure 6.**
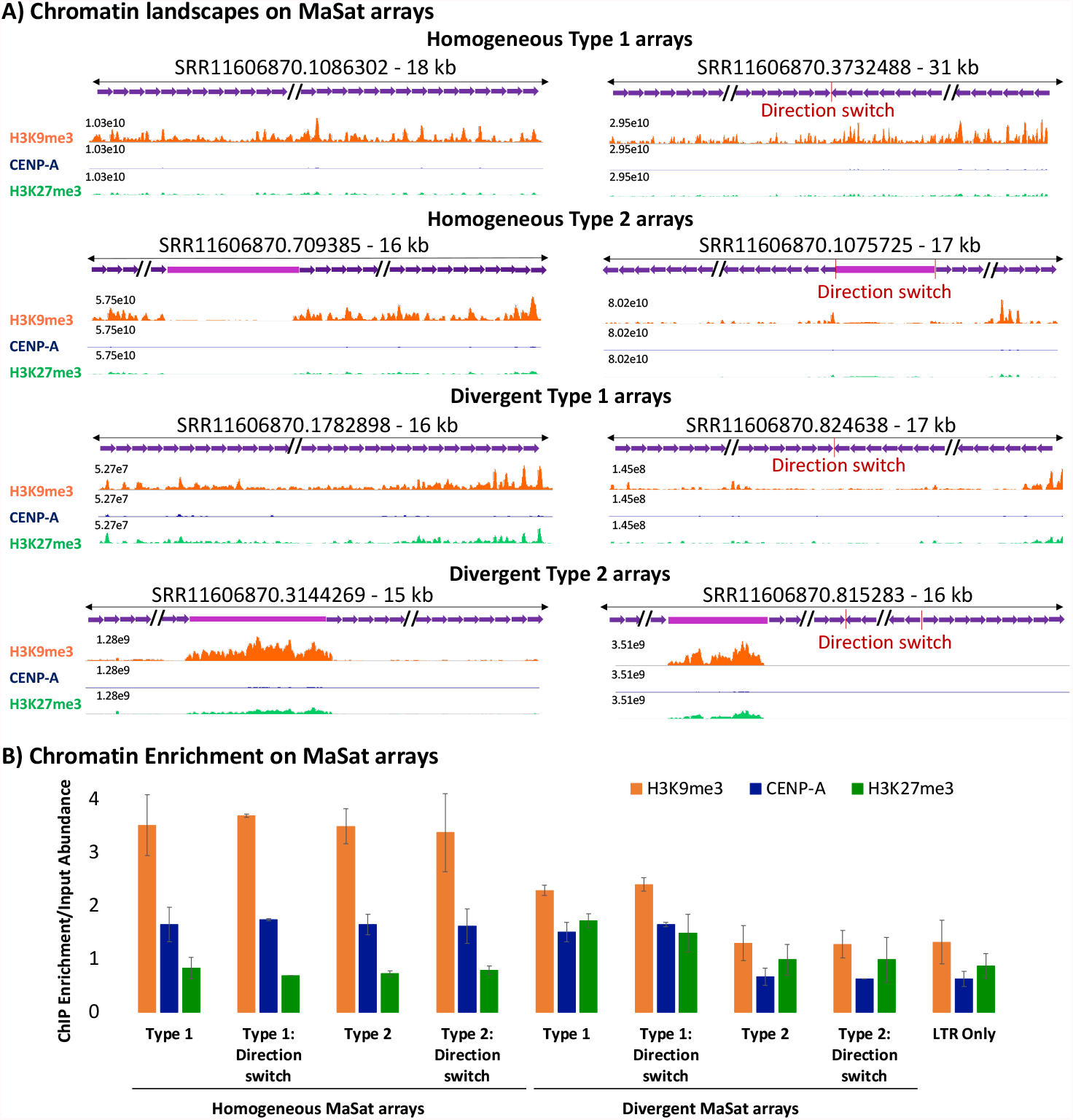
Occupancy of chromatin marks on pericentric MaSat arrays. **A)** CENP-A, H3K9me3, and H3K27me3 chromatin profiles on representative homogeneous and divergent MaSat arrays. The Y-axis is set to a constant value for a given array for CENP-A, H3K9me3, and H3K27me3 tracks. The length of each array is specified. The X-axis is not to the scale. **B)** CENP-A, H3K9me3, and H3K27me3 enrichment normalized to the satellite sequence abundance. The ChIP Enrichment was normalized to the input abundance and the normalized enrichment was averaged over three or more arrays for each type.

### CENP-A and H3K9me3 nucleosomes exhibit distinct organizations and conformations

Next, we analyzed the conformations of CENP-A and H3K9me3 containing nucleosomes on MiSat and MaSat arrays (Figure 7A, 7B). CENP-A chromatin displayed a general lack of nucleosome phasing on MiSat repeat units across all array types (Figure 7A). CENP-A peaks spanned either a single or multiple tandem MiSats suggesting that mouse CENP-A nucleosomes are tightly associated with other centromeric proteins to form larger complexes similar to those observed on human centromeric satellite arrays [28, 29]. Furthermore, CENP-A nucleosomes were present in several different conformations without any discernible arrangement relative to the CENP-B box. The variation in CENP-A conformation could not be related to the presence of an intact CENP-B box or its variants (Figure 7A). In contrast, H3K9me3 chromatin on MaSat satellites displayed a relatively uniform conformation with each peak occupying 234 bp MaSat unit (Figure 7B). Interestingly, although CENP-A and H3K27me3 chromatin were enriched only slightly on MaSats, they displayed a well-phased conformation, suggesting that homogenous MaSats exhibit the inherent property of phasing all types of chromatin (Figure 7B). Phasing observed in H3K9me3 nucleosomes on MaSats was absent on non-satellite sequences interspersed with MaSat. Divergent MaSat arrays exhibited less H3K9me3 phasing as compared to homogenous MaSat arrays (Figure 7B). Overall, CENP-A and H3K9me3 nucleosomes have different conformations suggesting distinct mechanisms of chromatin assembly at centromeric and pericentromeric regions.

**Figure 7.**
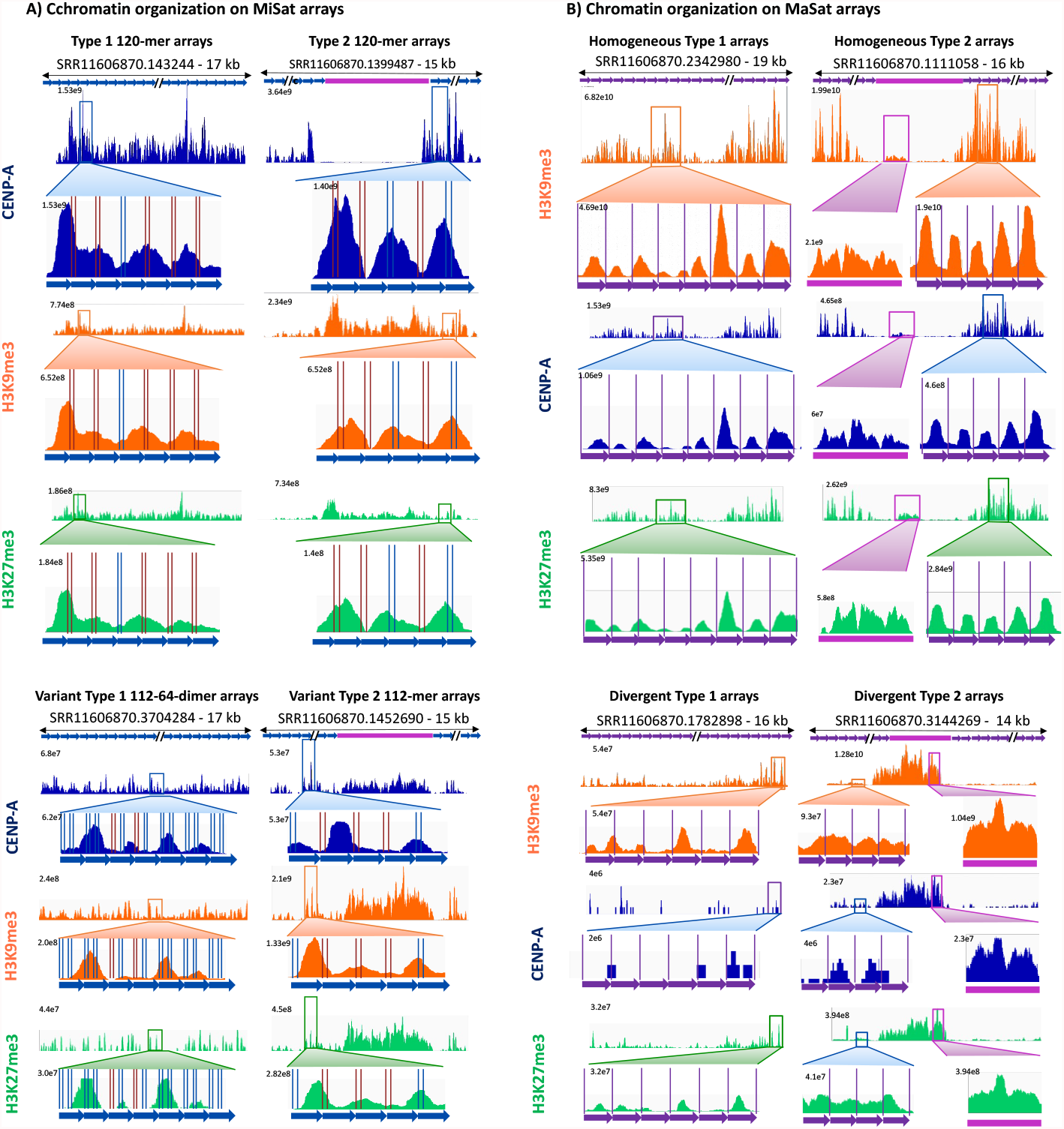
Distinct conformations of CENP-A, H3K9me3, and H3K27me3 containing nucleosomes. **A)** CENP-A, H3K9me3, and H3K27me3 ChIP-seq profiles on representative homogeneous and variant Type 1 and Type 2 MiSat arrays. **B)** CENP-A, H3K9me3, and H3K27me3 ChIP-seq profiles on representative homogeneous and divergent Type 1 and Type 2 MaSat arrays. CENP-B boxes on MiSat are marked by vertical blue (intact) and red lines (variant). MaSat monomers are separated by purple lines.

**Figure 8.**
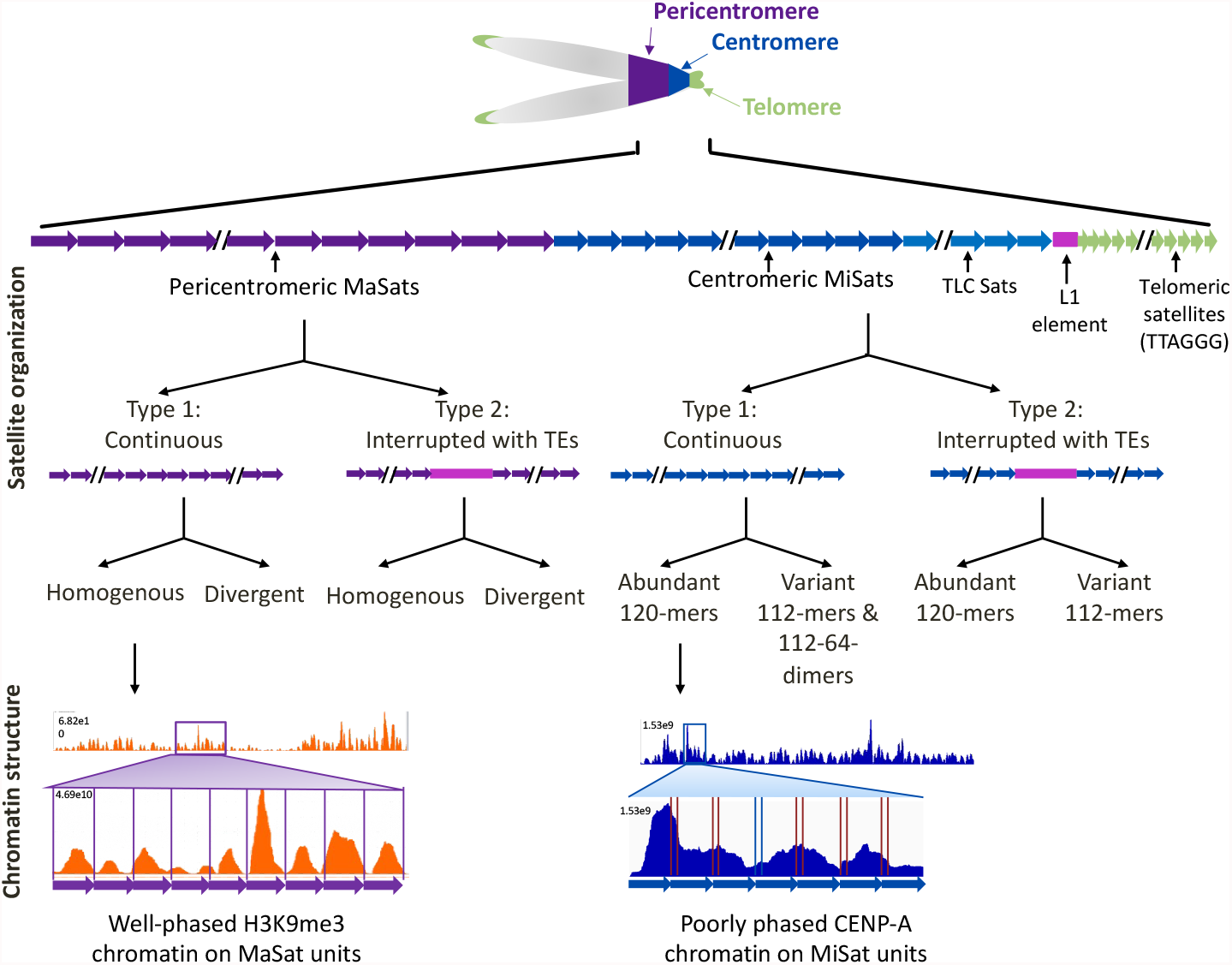
Schematic describing satellite and chromatin organization at mouse centromeres and pericentromeres.

## DISCUSSION

Previous studies have suggested that mouse centromeric MiSats and pericentromeric MaSats of autosomes and the X chromosome are highly homogeneous, with little variation within and across arrays [46, 47]. However, our results reveal significant variation between repeat units within and across major and minor satellite arrays (Figure 1-4, 8). For MiSat arrays, nucleotide variations were concentrated at the CENP-B box, including changes that can disrupt the ability of CENP-B to bind the CENP-B box [59]. The presence of CENP-B box variants at the house mouse centromeres suggests that variant CENP-B boxes might be associated with the differential binding of centromeric proteins on different centromeric satellite arrays. While CENP-B was initially considered not essential for centromere function due to the viability of CENP-B knockout mice and the absence of the CENP-B box on neocentromeres and Y centromeres [61, 65], recent studies have shown that lower CENP-B levels are associated with higher missegregation rates and lower fertility, suggesting that CENP-B plays an important role in centromere function and maintenance [62, 66, 67]. Furthermore, the CENP-B box density is correlated with the binding of CENP-A, CENP-B, and CENP-C at human centromeric chromatin [30].

In addition to the sequence variation among satellites within and across arrays, we also identified the presence of TEs at house mouse centromeres and pericentromeric regions. This finding is consistent with a previous study that identified TEs, specifically LINE and IAP elements, associated with MiSat [41]. TEs have also been previously identified in centromeric and pericentromeric regions of plants and some eukaryotes, including in humans and *Drosophila* [13, 68-70]. In humans, TEs are predominantly found in the pericentric region [9, 12]. In contrast, in *Drosophila*, islands of retroelements have been found at the functional regions of centromeres that bind CENP-A [13]. The role of TEs at centromeres is not well understood. Some studies have proposed that TEs are drivers of centromere evolution [68, 69]. The formation of new satellites from TE insertions at centromeres offers a potential explanation for the observed rapid evolution of centromeric sequences between species [68-70]. Centromeres have also been proposed to be genomic “safe” insertion zones for TEs, as surrounding repeats can act as a buffer [68, 69]. It has also been speculated that centromeric TEs are transcribed to non-coding RNAs that facilitate CENP-A deposition [69]. While we have identified long stretches of TEs interrupting mouse centromeres similar to *Drosophila*, these TEs do not bind CENP-A themselves. Instead, TEs at mouse centromeres are bound by a low level of repressive constitutive H3K9me3 heterochromatin, suggesting that they are kept in a somewhat silent state to avoid abnormally high transposon activity. Future functional studies will help understand the role of TEs at mouse centromeres.

Our findings of a surprising level of diversity in sequence and organization of mouse centromeric satellites within and across arrays indicate a potential conserved pattern of centromeric satellite variations between mice and humans. Although the extent of variations found in human centromere regions is higher compared to mice [9], our findings raise the possibility that mouse genomes may also contain chromosome-specific centromeric satellite arrays. Future studies using cytological analysis techniques will provide insight into the presence of chromosome-specific arrays at mouse centromeres. Additionally, our study highlights the variation in centromeric chromatin structure, even within a single MiSat array as previously seen in humans [30]. The differences in CENP-A organization between adjacent satellite units suggest that small sequence variations might affect the binding of CENP-A. Thus, CENP-A organization and binding in the house mouse may have a sequence-dependent component. Overall, our findings on the sequence and organization of mouse centromeric satellite and chromatin shed light on the dynamic yet conserved pattern of satellite sequences and organization and provide a basis for future studies on the functional implications of centromeric satellite diversity in mammals.

## METHODS

### Animals and tissue homogenization

The C57BL/6 strain was purchased from the Jackson Laboratory and maintained following the institutional animal care and use committee guidelines. Liver tissues from euthanized C57BL/6 were snap-frozen in liquid nitrogen and ground to powder using a mortar and pestle. The powder was resuspended in 1x PBS containing Roche protease inhibitor cocktail (Millipore Sigma Cat# 11836170001) and dounced with a 15 ml glass douncer using 50 strokes on ice. Glass dounced homogenate was further homogenized using the Tekmar homogenizer on ice. The resulting suspension was passed through a 50-micron nylon filter, the flow through was pelleted at 1800 rpm at 4°C, and the pellet was washed with 1X PBS. The pellet containing homogenized cells was resuspended in 1X PBS.

### Chromatin Immunoprecipitation (ChIP)

Native ChIP was performed using the protocol described previously [29] on homogenized liver cells using anti-CENP-A (Cell Signaling Technologies, Cat # C51A7), H3K9me3 (Abcam, Cat # ab8898), H3K27me3 (Cell Signaling Technologies, Cat # C36B11) and IgG (Abcam, Cat # ab46540) antibodies using the following modification. After MNase digestion and needle extraction, low (150 mM) and high (350 mM) NaCl containing fractions were combined for antibody binding and downstream steps.

### Analysis of long-read HiFi sequencing data

We analyzed long-read HiFi sequencing data generated using the PacBio Sequel II system for C57BL/6J mouse genome from Hon et al. (2020) [50]. The read length distribution of all reads was calculated using BBMap global aligner from the Joint Genome Institute [71]. To isolate major and minor satellite arrays, the LRS data was searched against libraries of *Mus musculus* major and minor satellite reference sequences using NCBI BLAST. The read length distributions of reads with minor and major satellites were calculated using BBMap. The long reads identified to contain at least one minor or major satellite were then further searched against the RepeatMasker database to characterize TEs and other repeats in the arrays (RepeatMasker -species “Mus musculus” -a). MiSat 112-mer and 112-64-mer variants were identified by searching MiSat long reads against reference sequences [49] using NCBI BLAST. MaSat arrays containing > 10 repeats with less than 75% sequence similarity to *Mus musculus* major satellite reference sequence were classified as Divergent MaSat arrays. CENP-B box sequences from all minor satellites were extracted and clustered using CD-HIT to analyze CENP-B box variants [72, 73]. Sample satellite containing arrays (36 minor, 6 centromere-telomere junctions, and 39 major) identified from the LRS data were selected for further analysis.

### Assembly of HiFi sequencing reads

We created predicted contigs from the PacBio long reads using Hifiasm default parameters [64]. Next, we isolated the contigs containing MiSat and MaSat using NCBI BLAST. We removed any contigs with less than 100 satellites from further analyses. We obtained 18 MiSat contigs and 138 MaSat contigs. Only the top 50 MaSat contigs were analyzed to determine the arrangement of satellites.

### Library preparation, and sequencing

Libraries were prepared from ChIP DNA fragments using the KAPA HyperPrep Kit following the KAPA HyperPrep Kit manual using 12 library amplification cycles. The library was sequenced using the NextSeq 500/550 Mid Output Kit with approximately 9.3 million paired-end 75 bp reads per sample.

### Analysis of chromatin profiling data

The sequencing reads from CENP-A, H3K9me3, H3K27me3, and IgG ChIP sequencing were mapped to sample minor and major satellite containing LRS reads using Bowtie2 (bowtie2 --end-to-end --very-sensitive --no-mixed --no-discordant -q --phred33 -I 10 -X 700) [74].The sam files generated by Bowtie2 were converted to bed files using samtools and bedtools. The bedgraphs were generated using a custom script and visualized on the Integrated Genome Viewer (IGV) [75].

## Funding

This work was supported by 1R35GM147558-01 NIH grant to JT. JP was funded by a Summer Undergraduate Research Experience fellowship by College of Arts and Sciences at Emory University in 2021.

## Acknowledgment

Sequencing of ChIP-Seq samples was performed at Emory Integrated Genomics Core. We thank William Kelly, Roger Deal, David Katz, and Thakur lab members for their feedback on the manuscript.

## Conflict of interest

Authors report no conflict of interest.

## Notes

### Competing Interest Statement

The authors have declared no competing interest.

## REFERENCES

1. Thompson SL, Bakhoum SF, Compton DA: Mechanisms of chromosomal instability. Curr Biol 2010, 20: R285–295.

2. Ting DT, Lipson D, Paul S, Brannigan BW, Akhavanfard S, Coffman EJ, Contino G, Deshpande V, Iafrate AJ, Letovsky S, et al: Aberrant overexpression of satellite repeats in pancreatic and other epithelial cancers. Science 2011, 331: 593–596.

3. Barra V, Fachinetti D: The dark side of centromeres: types, causes and consequences of structural abnormalities implicating centromeric DNA. Nature Communications 2018, 9: 4340–4340.

4. Gemble S, Simon A, Pennetier C, Dumont M, Herve S, Meitinger F, Oegema K, Rodriguez R, Almouzni G, Fachinetti D, Basto R: Centromere Dysfunction Compromises Mitotic Spindle Pole Integrity. Curr Biol 2019, 29: 3072–3080 e3075.

5. McKinley KL, Cheeseman IM: The molecular basis for centromere identity and function. Nature Reviews Molecular Cell Biology 2016, 17: 16–29.

6. Palmer D, O’Day K, Wener M, Andrews B, Margolis R: A 17-kD centromere protein (CENP-A) copurifies with nucleosome core particles and with histones. The Journal of Cell Biology 1987, 104: 805–815.

7. Thakur J, Packiaraj J, Henikoff S: Sequence, Chromatin and Evolution of Satellite DNA. Int J Mol Sci 2021, 22.

8. Rudd MK, Willard HF: Analysis of the centromeric regions of the human genome assembly. Trends Genet 2004, 20: 529–533.

9. Altemose N, Logsdon GA, Bzikadze AV, Sidhwani P, Langley SA, Caldas GV, Hoyt SJ, Uralsky L, Ryabov FD, Shew CJ, et al: Complete genomic and epigenetic maps of human centromeres. Science 2022, 376.

10. Schueler MG, Higgins AW, Rudd MK, Gustashaw K, Willard HF: Genomic and genetic definition of a functional human centromere. Science 2001, 294: 109–115.

11. Alexandrov I, Kazakov A, Tumeneva I, Shepelev V, Yurov Y: Alpha-satellite DNA of primates: old and new families. Chromosoma 2001, 110: 253–266.

12. Waye JS, Willard HF: Human beta satellite DNA: genomic organization and sequence definition of a class of highly repetitive tandem DNA. Proc Natl Acad Sci U S A 1989, 86: 6250–6254.

13. Chang CH, Chavan A, Palladino J, Wei X, Martins NMC, Santinello B, Chen CC, Erceg J, Beliveau BJ, Wu CT, et al: Islands of retroelements are major components of Drosophila centromeres. PLoS Biology 2019, 17.

14. Alexandrov IA, Medvedev LI, Mashkova TD, Kisselev LL, Romanova LY, Yurov YB: Definition of a new alpha satellite suprachromosomal family characterized by monomeric organization. Nucleic Acids Res 1993, 21: 2209–2215.

15. Melters DP, Bradnam KR, Young HA, Telis N, May MR, Ruby JG, Sebra R, Peluso P, Eid J, Rank D, et al: Comparative analysis of tandem repeats from hundreds of species reveals unique insights into centromere evolution. Genome Biology 2013, 14: 1–20.

16. Plohl M, Meštrović N, Mravinac B: Centromere identity from the DNA point of view. Chromosoma 2014, 123: 313–325.

17. Smith OK, Limouse C, Fryer KA, Teran NA, Sundararajan K, Heald R, Straight AF: Identification and characterization of centromeric sequences in Xenopus laevis. Genome Res 2021, 31: 958–967.

18. Padmanabhan S, Thakur J, Siddharthan R, Sanyal K: Rapid evolution of Cse4p-rich centromeric DNA sequences in closely related pathogenic yeasts, Candida albicans and Candida dubliniensis. Proc Natl Acad Sci U S A 2008, 105: 19797–19802.

19. Voullaire LE, Slater HR, Petrovic V, Choo KHA: A Functional Marker Centromere with No Detectable Alpha-Satellite, Satellite Ill, or CENP-B Protein: Activation of a Latent Centromere? Am J Hum Genet 1993, 52: 1153–1163.

20. Heun P, Erhardt S, Blower MD, Weiss S, Skora AD, Karpen GH: Mislocalization of the drosophila centromere-specific histone CID promotes formation of functional ectopic kinetochores. Developmental Cell 2006, 10: 303–315.

21. Thakur J, Sanyal K: Efficient neocentromere formation is suppressed by gene conversion to maintain centromere function at native physical chromosomal loci in Candida albicans. Genome Res 2013, 23: 638–652.

22. Morrison O, Thakur J: Molecular complexes at euchromatin, heterochromatin and centromeric chromatin. International Journal of Molecular Sciences 2021, 22: 6922.

23. Dalal Y, Wang H, Lindsay S, Henikoff S: Tetrameric structure of centromeric nucleosomes in interphase Drosophila cells. PLoS biology 2007, 5: e218.

24. Dimitriadis EK, Weber C, Gill RK, Diekmann S, Dalal Y: Tetrameric organization of vertebrate centromeric nucleosomes. Proc Natl Acad Sci U S A 2010, 107: 20317–20322.

25. Krassovsky K, Henikoff JG, Henikoff S: Tripartite organization of centromeric chromatin in budding yeast. Proc Natl Acad Sci U S A 2012, 109: 243–248.

26. Hasson D, Panchenko T, Salimian KJ, Salman MU, Sekulic N, Alonso A, Warburton PE, Black BE: The octamer is the major form of CENP-A nucleosomes at human centromeres. Nature structural & molecular biology 2013, 20: 687–687.

27. Westhorpe FG, Straight AF: The centromere: epigenetic control of chromosome segregation during mitosis. Cold Spring Harb Perspect Biol 2014, 7: a015818.

28. Henikoff JG, Thakur J, Kasinathan S, Henikoff S: A unique chromatin complex occupies young a-satellite arrays of human centromeres. Science Advances 2015, 1.

29. Thakur J, Henikoff S: CENPT bridges adjacent CENPA nucleosomes on young human alpha-satellite dimers. Genome Res 2016, 26: 1178–1187.

30. Thakur J, Henikoff S: Unexpected conformational variations of the human centromeric chromatin complex. Genes and Development 2018, 32: 20–25.

31. Peters AH, Kubicek S, Mechtler K, O’Sullivan RJ, Derijck AA, Perez-Burgos L, Kohlmaier A, Opravil S, Tachibana M, Shinkai Y, et al: Partitioning and plasticity of repressive histone methylation states in mammalian chromatin. Mol Cell 2003, 12: 1577–1589.

32. Rice JC, Briggs SD, Ueberheide B, Barber CM, Shabanowitz J, Hunt DF, Shinkai Y, Allis CD: Histone methyltransferases direct different degrees of methylation to define distinct chromatin domains. Mol Cell 2003, 12: 1591–1598.

33. Rosenfeld JA, Wang Z, Schones DE, Zhao K, DeSalle R, Zhang MQ: Determination of enriched histone modifications in non-genic portions of the human genome. BMC Genomics 2009, 10: 143.

34. Bernard P, Maure J-F, Partridge JF, Genier S, Javerzat J-P, Allshire RC: Requirement of heterochromatin for cohesion at centromeres. Science 2001, 294: 2539–2542.

35. Peters AH, O’Carroll D, Scherthan H, Mechtler K, Sauer S, Schofer C, Weipoltshammer K, Pagani M, Lachner M, Kohlmaier A, et al: Loss of the Suv39h histone methyltransferases impairs mammalian heterochromatin and genome stability. Cell 2001, 107: 323–337.

36. Treangen TJ, Salzberg SL: Repetitive DNA and next-generation sequencing: computational challenges and solutions. Nat Rev Genet 2011, 13: 36–46.

37. Wenger AM, Peluso P, Rowell WJ, Chang P-C, Hall RJ, Concepcion GT, Ebler J, Fungtammasan A, Kolesnikov A, Olson ND, et al: Accurate circular consensus long-read sequencing improves variant detection and assembly of a human genome. Nature Biotechnology 2019, 37: 1155–1162.

38. Logsdon GA, Vollger MR, Eichler EE: Long-read human genome sequencing and its applications. Nat Rev Genet 2020, 21: 597–614.

39. Nurk S, Koren S, Rhie A, Rautiainen M, Bzikadze AV, Mikheenko A, Vollger MR, Altemose N, Uralsky L, Gershman A, et al: The complete sequence of a human genome. Science 2022, 376: 44–53.

40. Kipling D, Ackford HE, Taylor BA, Cooke HJ: Mouse minor satellite DNA genetically maps to the centromere and is physically linked to the proximal telomere. Genomics 1991, 11: 235–241.

41. Komissarov AS, Kuznetsova IS, Podgornaia OI: Mouse centromeric tandem repeats in silico and in situ. Genetika 2010, 46: 1217–1221.

42. Komissarov AS, Gavrilova EV, Demin SJ, Ishov AM, Podgornaya OI: Tandemly repeated DNA families in the mouse genome. BMC Genomics 2011, 12: 531.

43. Guenatri M, Bailly D, Maison C, Almouzni G: Mouse centric and pericentric satellite repeats form distinct functional heterochromatin. Journal of Cell Biology 2004, 166: 493–505.

44. Iwata-Otsubo A, Dawicki-McKenna JM, Akera T, Falk SJ, Chmátal L, Yang K, Sullivan BA, Schultz RM, Lampson MA, Black BE: Expanded Satellite Repeats Amplify a Discrete CENP-A Nucleosome Assembly Site on Chromosomes that Drive in Female Meiosis. Current Biology 2017, 27: 2365-2373.e2368.

45. Almouzni G, Probst AV: Heterochromatin maintenance and establishment: lessons from the mouse pericentromere. Nucleus 2011, 2: 332–338.

46. Kalitsis P, Griffiths B, Choo KHA: Mouse telocentric sequences reveal a high rate of homogenization and possible role in Robertsonian translocation. Proceedings of the National Academy of Sciences of the United States of America 2006, 103: 8786–8791.

47. Wong AK, Rattner JB: Sequence organization and cytological localization of the minor satellite of mouse. Nucleic Acids Res 1988, 16: 11645–11661.

48. Arora UP, Charlebois C, Lawal RA, Dumont BL: Population and subspecies diversity at mouse centromere satellites. BMC Genomics 2021, 22.

49. Rice WR: Centromeric repeats of the Western European house mouse I: high sequence diversity among monomers at local and global spatial scales. Cold Spring Harbor Laboratory; 2020.

50. Hon T, Mars K, Young G, Tsai YC, Karalius JW, Landolin JM, Maurer N, Kudrna D, Hardigan MA, Steiner CC, et al: Highly accurate long-read HiFi sequencing data for five complex genomes. Sci Data 2020, 7: 399.

51. Sharif J, Shinkai Y, Koseki H: Is there a role for endogenous retroviruses to mediate long-term adaptive phenotypic response upon environmental inputs? Philosophical Transactions of the Royal Society B: Biological Sciences 2013, 368.

52. Kuff EL, Lueders KK: The Intracisternal A-Particle Gene Family: Structure and Functional Aspects. Advances in Cancer Research 1988, 51: 183–276.

53. Qin C, Wang Z, Shang J, Bekkari K, Liu R, Pacchione S, McNulty KA, Ng A, Barnum JE, Storer RD: Intracisternal a particle genes: Distribution in the mouse genome, active subtypes, and potential roles as species-specific mediators of susceptibility to cancer. Molecular Carcinogenesis 2010, 49: 54–67.

54. Lunyak VV, Prefontaine GG, Nunez E, Cramer T, Ju BG, Ohgi KA, Hutt K, Roy R, Garcia-Diaz A, Zhu X, et al: Developmentally regulated activation of a SINE B2 repeat as a domain boundary in organogenesis. Science 2007, 317: 248–251.

55. Bourque G, Leong B, Vega VB, Chen X, Lee YL, Srinivasan KG, Chew JL, Ruan Y, Wei CL, Ng HH, Liu ET: Evolution of the mammalian transcription factor binding repertoire via transposable elements. Genome Res 2008, 18: 1752–1762.

56. Schmidt D, Schwalie PC, Wilson MD, Ballester B, Goncalves A, Kutter C, Brown GD, Marshall A, Flicek P, Odom DT: Waves of retrotransposon expansion remodel genome organization and CTCF binding in multiple mammalian lineages. Cell 2012, 148: 335–348.

57. Lander ES, Linton LM, Birren B, Nusbaum C, Zody MC, Baldwin J, Devon K, Dewar K, Doyle M, FitzHugh W, et al: Initial sequencing and analysis of the human genome. Nature 2001, 409: 860–921.

58. Masumoto H, Masukata H, Muro Y, Nozaki N, Okazaki T: A human centromere antigen (CENP-B) interacts with a short specific sequence in alphoid DNA, a human centromeric satellite. Journal of Cell Biology 1989, 109: 1963–1973.

59. Masumoto H, Yoda K, Ikeno M, Kitagawa K, Muro Y, Okazaki T: Properties of CENP-B and its target sequence in a satellite DNA. In Chromosome segregation and aneuploidy. Springer; 1993: 31–43.

60. Tanaka Y, Nureki O, Kurumizaka H, Fukai S, Kawaguchi S, Ikuta M, Iwahara J, Okazaki T, Yokoyama S: Crystal structure of the CENP-B protein-DNA complex: the DNA-binding domains of CENP-B induce kinks in the CENP-B box DNA. EMBO J 2001, 20: 6612–6618.

61. Kapoor M, Montes De Oca Luna R, Liu G, Lozano G, Cummings C, Mancini M, Ouspenski I, Brinkley BR, May GS: The cenpB gene is not essential in mice. Chromosoma 1998, 107: 570–576.

62. Fachinetti D, Han JS, McMahon MA, Ly P, Abdullah A, Wong AJ, Cleveland DW: DNA Sequence-Specific Binding of CENP-B Enhances the Fidelity of Human Centromere Function. Developmental Cell 2015, 33: 314–327.

63. Joseph A, Mitchell AR, Miller OJ: The organization of the mouse satellite DNA at centromeres. Exp Cell Res 1989, 183: 494–500.

64. Cheng H, Concepcion GT, Feng X, Zhang H, Li H: Haplotype-resolved de novo assembly using phased assembly graphs with hifiasm. Nat Methods 2021, 18: 170–175.

65. Depinet TW, Zackowski JL, Earnshaw WC, Kaffe S, Sekhon GS, Stallard R, Sullivan BA, Vance GH, Van Dyke DL, Willard HF, et al: Characterization of neo-centromeres in marker chromosomes lacking detectable alpha-satellite DNA. Hum Mol Genet 1997, 6: 1195–1204.

66. Morozov VM, Giovinazzi S, Ishov AM: CENP-B protects centromere chromatin integrity by facilitating histone deposition via the H3.3-specific chaperone Daxx. Epigenetics Chromatin 2017, 10: 63.

67. Chardon F, Japaridze A, Witt H, Velikovsky L, Chakraborty C, Wilhelm T, Dumont M, Yang W, Kikuti C, Gangnard S, et al: CENP-B-mediated DNA loops regulate activity and stability of human centromeres. Molecular Cell 2022, 82: 1751-1767.e1758.

68. Wong LH, Choo KA: Evolutionary dynamics of transposable elements at the centromere. TRENDS in Genetics 2004, 20: 611–616.

69. Klein SJ, O’Neill RJ, Klein SJ, O’Neill RJ: Transposable elements: genome innovation, chromosome diversity, and centromere conflict. Chromosome Research 2018 26:1 2018, 26: 5–23.

70. Meštrović N, Mravinac B, Pavlek M, Vojvoda-Zeljko T, Šatović E, Plohl M: Structural and functional liaisons between transposable elements and satellite DNAs. Chromosome Res 2015, 23: 583–596.

71. Bushnell B: BBMap: A Fast, Accurate, Splice-Aware Aligner. Lawrence Berkeley National Lab 2014.

72. Fu L, Niu B, Zhu Z, Wu S, Li W: CD-HIT: accelerated for clustering the next-generation sequencing data. Bioinformatics 2012, 28: 3150–3152.

73. Li W, Godzik A: Cd-hit: A fast program for clustering and comparing large sets of protein or nucleotide sequences. Bioinformatics 2006, 22: 1658–1659.

74. Langmead B, Salzberg SL: Fast gapped-read alignment with Bowtie 2. Nat Methods 2012, 9: 357–359.

75. Robinson JT, Thorvaldsdottir H, Winckler W, Guttman M, Lander ES, Getz G, Mesirov JP: Integrative genomics viewer. Nat Biotechnol 2011, 29: 24–26.

