## Supplementary Figures for "DNA satellite and chromatin organization at house mouse centromeres and pericentromeres"

### A) Read length distribution

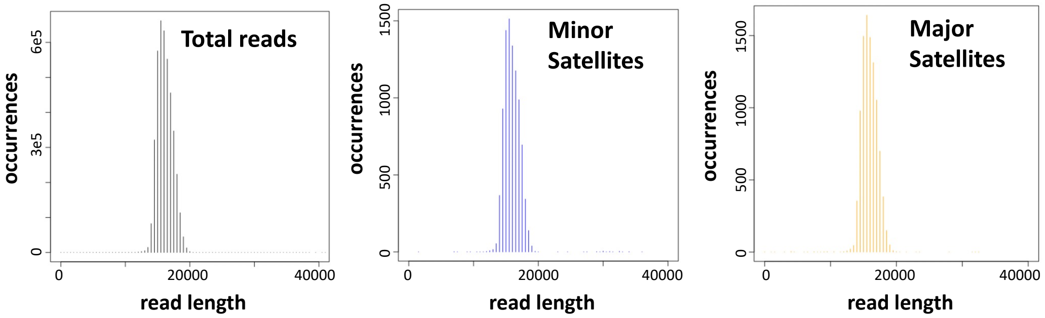


### B) MiSat arrays

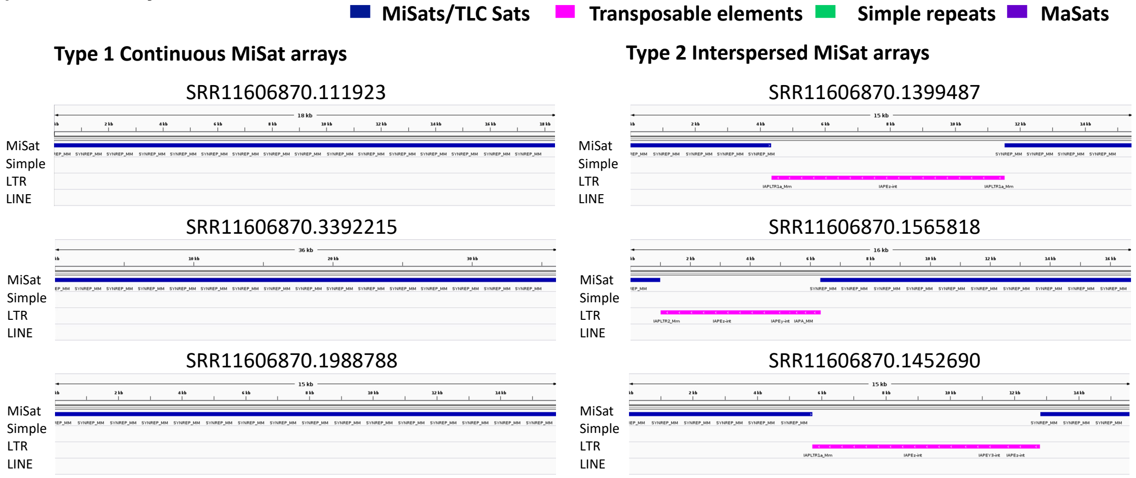


### C) Centromere and telomere junctions

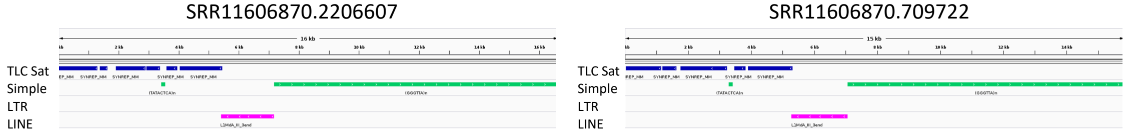


### D) MaSat arrays

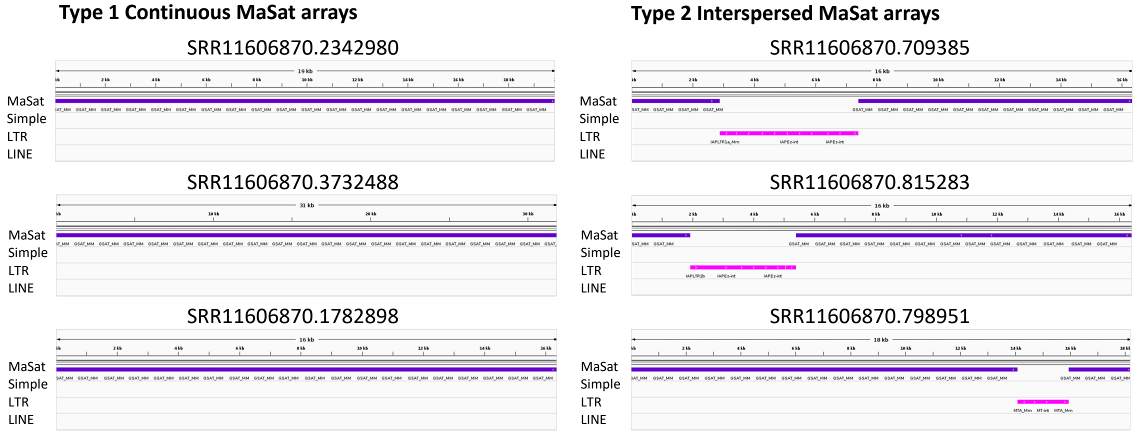

**Supplementary Figure 1. A) Read length distribution in the LRS data analyzed in this study.** Detailed organization of sample LRS reads with B) MiSat arrays, C) Centromere-Telomere junctions, and D) and MaSat arrays.

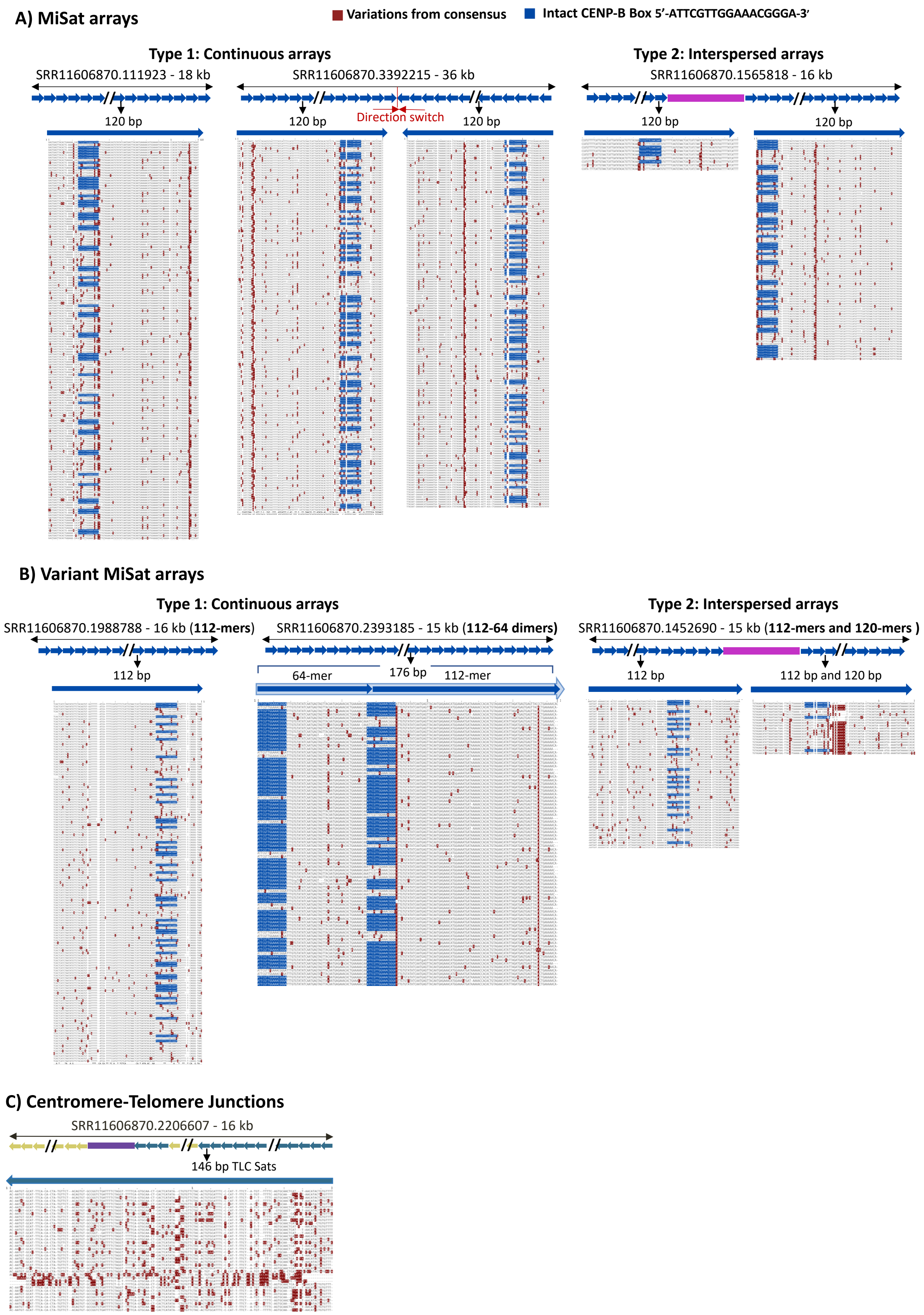

A) MiSat arrays
Variations from consensus
Intact CENP-B Box 5'-ATTCGTTGGAAACGGGA-3'
Type 1: Continuous arrays
SRR11606870.111923 - 18 kb
SRR11606870.3392215 - 36 kb
120 bp
120 bp
Direction switch
120 bp
Type 2: Interspersed arrays
SRR11606870.1565818 - 16 kb
120 bp
120 bp
B) Variant MiSat arrays
Type 1: Continuous arrays
SRR11606870.1988788 - 16 kb (112-mers)
SRR11606870.2393185 - 15 kb (112-64 dimers)
112 bp
64-mer
176 bp
112-mer
Type 2: Interspersed arrays
SRR11606870.1452690 - 15 kb (112-mers and 120-mers )
112 bp
112 bp and 120 bp
C) Centromere-Telomere Junctions
SRR11606870.2206607 - 16 kb
146 bp TLC Sats

### Supplementary Figure 2. Full alignments from MiSat and telomere-centromere junction TLC arrays.

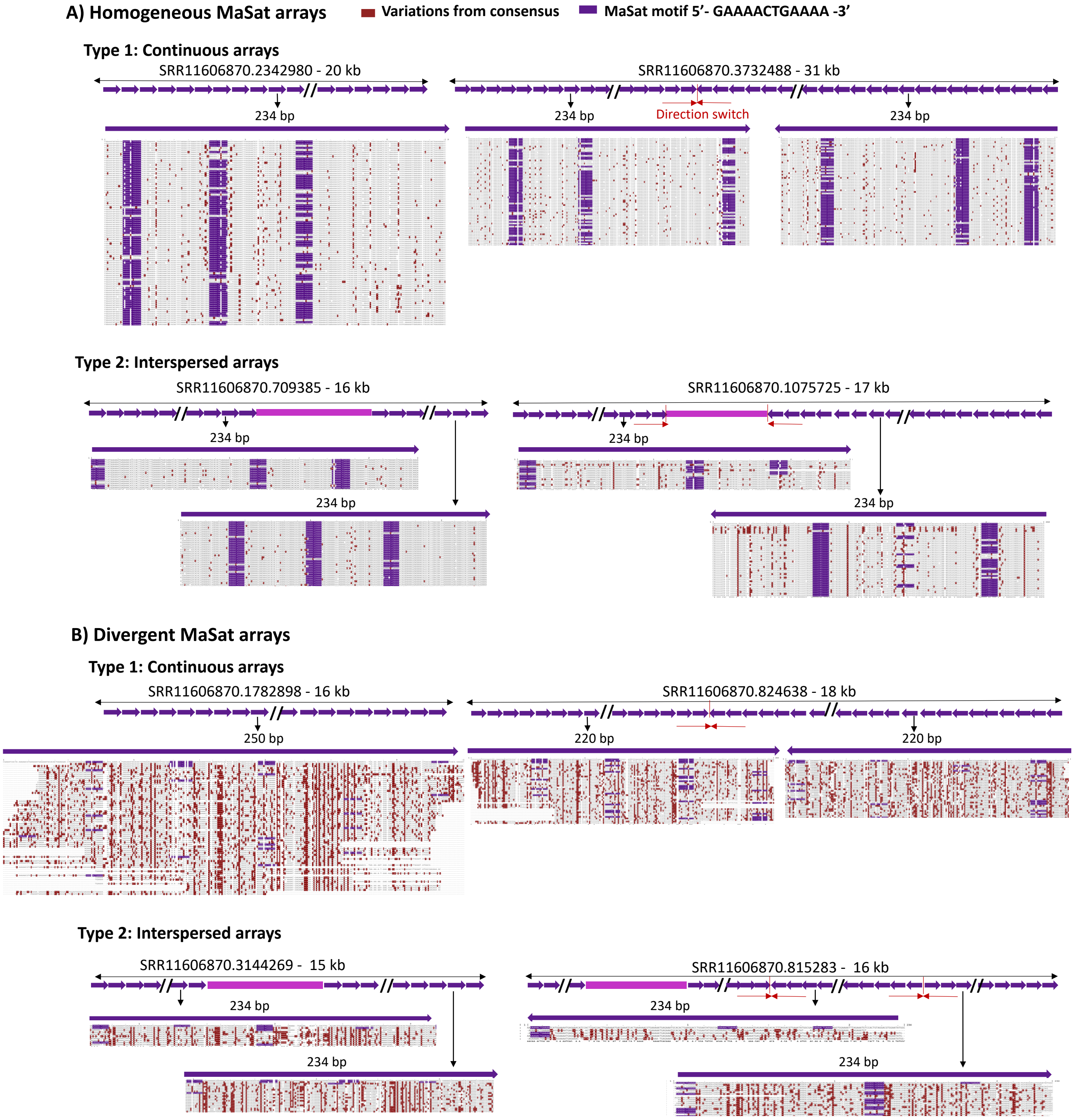


### Supplementary Figure 3. Full alignments from MaSat arrays.

### Chromatin landscape at centromere-telomere junction arrays

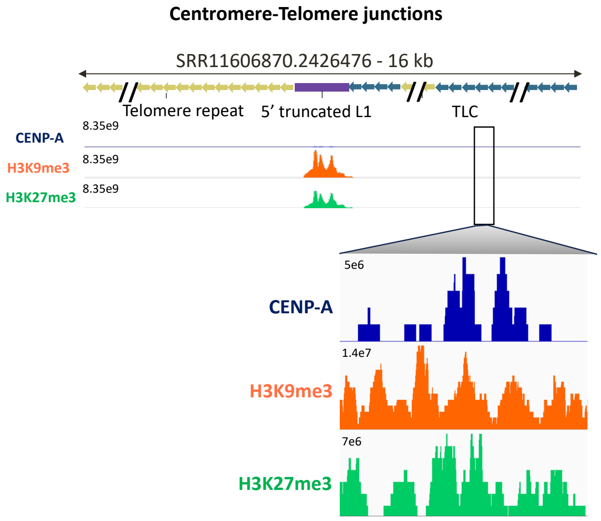


**Supplementary Figure 4. CENP-A, H3K9me3, and H3K27me3 chromatin profiles at centromere-telomere junction arrays including TLC arrays.**
